# Host cell factors important for BHV-1 cell entry revealed by genome-wide CRISPR knockout screen

**DOI:** 10.1101/2020.06.18.160523

**Authors:** Wenfang Spring Tan, Enguang Rong, Inga Dry, Simon Lillico, Andy Law, Bruce Whitelaw, Robert G. Dalziel

## Abstract

In order to identify host factors that impact Bovine Herpes Virus Type 1 (BHV-1) infection we previously applied a genome wide CRISPR knockout screen with a library covering all bovine protein coding genes. We compiled a list of both pro-viral and anti-viral proteins involved in BHV-1 replication; here we provide further analysis of those that are potentially involved in viral entry into the host cell. These entry related factors include the cell surface proteins PVR and PVRL2, a group of enzymes directly or indirectly associated with the biosynthesis of Heparan Sulfate Proteoglycans (HSPG), and proteins that reside in the Golgi apparatus engaging in intra-Golgi trafficking. For the first time, we provide evidence that PVRL2 serves a receptor for BHV-1, mediating more efficient entry than the previously identified PVR. By knocking out two enzymes that catalyze HSPG chain elongation, HST2ST1 and GLCE, we demonstrated the significance of HSPG in BHV-1 entry. Another intriguing cluster of genes, COG1, COG2 and COG4-7 encodes for six subunits of the conserved oligomeric Golgi (COG) complex. MDBK cells lacking COG6 were less infectable by BHV-1 but release newly produced virions more efficiently as evidenced by fewer but bigger plaques compared to control cells, suggesting impaired HSPG biosynthesis. To facilitate candidate validation, we devised a one-step multiplex CRISPR interference (CRISPRi) system named CRISPR3i that enables quick and simultaneous deployment of three CRISPRs for efficient gene inactivation. Using CRISPR3i, we verified an additional 23 candidates, with many implicated in cellular entry.

## Introduction

Bovine Herpes Virus Type 1 (BHV-1) is a widespread virus. Acute infection or reactivation from latency causes severe syndromes of the respiratory and reproductive systems, transient immunosuppression and co-infection with other microbes, leading to substantial economic loss annually to cattle industries worldwide^1–3^. Like other alpha herpesviruses, BHV-1 is an enveloped large double strand DNA virus that enters the host cell via receptor binding mediated membrane fusion either on the cell surface or within endosomes^4^. BHV-1 entry is initiated by interaction between cellular surface molecules and viral envelope glycoproteins gB, gC, and gD^2,5,6^. Although not required for replication, gC plays a major role in attachment by interacting with cell surface heparan sulfate proteoglycan (HSPG) ^7,8^, thereby bringing gB and gD into proximity with their receptors^2,9–13^. Specifically, gD binds to cellular receptors nectin-1(PVRL1) and poliovirus receptor (PVR)^14–16^ while gB or a gH/gL complex may interact with putative alphaherpesvirus gB-receptor PILRα, resulting in fusion of the viral envelope with the cell membrane^17–22^. Previous studies found that human cell surface proteins nectin-2 or PVRL2 can mediate efficient entry of Herpes Simplex Virus 2 and mutant strains of Herpes Simplex Virus 1 (HSV-1) but not wild-type HSV-1 or BHV-1 into Chinese Hamster Ovary (CHO) cells^15,23,24^. Perhaps due to this negative outcome, the possibility of bovine nectin-2 serving as a receptor for BHV-1 entry into cow cells has never been further explored.

Ubiquitously expressed by most mammalian cell types, HSPG is primarily anchored in the plasma membranes or extracellular matrix. The transmembrane core protein of HSPG is covalently linked to chains of glycosaminoglycan (GAG) formed by unbranched sulfated polysaccharides, also known as heparan sulfates (HS)^25^. Other GAGs such as chondroitin sulfates (CS) and dermatan sulfates (DS) can also be attached to these core proteins^26^. The negatively charged GAG chains interact electrostatically with the basic residues of either the glycoproteins of enveloped viruses or the capsid proteins of non-enveloped viruses. A broad range of viruses, including Human Immunodeficiency Virus Type 1, Respiratory Syncytial Virus (RSV), Hepatitis E virus (HEV), HSV-1, Foot and Mouth Disease Virus^27^ and BHV-1^7,8^, overcome Brownian motion by exploiting such weak interactions and thereby increase their presence at the host cell surface, boosting the probability of binding to more specific entry receptors for internalization^25^. HSPG can even serve as a direct entry receptor for some of these viruses (HSV-1, HEV, and RSV)^28–30^ while other GAGs have also been implicated in viral entry into host cells^31^, e.g. CS binding by HSV-1^32,33^.

Like many other glycosylation and sulfation processes, the biosynthesis of HSPG is a multi-step reaction that takes place in the Golgi network. To initiate the process, biosynthetic precursors including 3′-phosphoadenosine-5′-phosphosulfate (PAPS) and UDP-sugars are transported into the Golgi lumen from the cytosol to serve as substrates^34^. Once the linkage region is attached to a serine residue in the core protein, the exostosins (EXT1, EXT2, EXTL1-3)^27,35^ add alternating units of N-Acetylglucosamine (GlcNAc) and glucuronic acid (GlcA) to the growing chain, followed by a series of sulfation and epimerization modifications to the units^26^. Upon completion, the core proteins decorated with these HS chains are transported towards the plasma membrane for exocytosis. The proper distribution of these exostosins is crucial for successful completion of HS biosynthesis, and this is maintained by retrograde vesicle traffic between Golgi cisternae^36^. Vesicle tethering at the Golgi apparatus prior to fusion is mediated by Rab GTPase coordinated interactions between the coiled-coil golgin tethers and the multi-subunit tethering complex known as the conserved oligomeric Golgi (COG) complex^36^. The COG complex is crucial for multiple aspects of the Golgi apparatus biology, including maintenance of its homeostasis, intracellular trafficking and protein glycosylation^37^. The significance of vesicle tethering in the context of glycan biosynthesis is highlighted by a set of diseases known as congenital disorders of glycosylation, caused by defects in COG subunits^38,39^. It has also been shown that COG perturbation in CHO cells leads to a defect in *O*-linked glycosylation^40,41^.

The recent development of robust genome editing tools enables us to interrogate host genes putatively associated with viral infections. These tools, such as the TAL effector nucleases (TALENs)^42,43^ and CRISPR/Cas9^44–46^, can be programmed to target almost any DNA sequence and induce a double strand break (DSB). In the absence of an exogenous DNA template the DSB is primarily repaired by non-homologous end joining (NHEJ), often leading to frameshifting indels and gene knockout (KO). CRISPR/Cas9 relies on base pairing between the 20-bp seed sequence of the guide and target DNA for specificity, a unique characteristic that enabled the development of CRISPR libraries targeting all protein coding genes in humans, mice and the cow, facilitating the study of genes pivotal to host-pathogen interactions in a high throughput manner^47–50^. As an alternative to gene knockout, Cas9 can also be repurposed to either upregulate or downregulate endogenous genes. To achieve this the two endonuclease domains of the Cas9 protein are deactivated to produce a catalytically “dead” Cas9 (dCas9), and gene transactivating or inactivating domains are fused to this to activate (CRISPRa)^51^ or interfere (CRISPRi)^52^ with gene transcription. For CRISPRi, a fusion complex consisting of dCas9, KRAB and the TRD domain of MeCP2 has proven to be efficient in gene silencing^52^. To ensure efficient down regulation of host gene expression, accurate Transcription Start Site (TSS) annotation is required as the guide RNA must be designed to bind immediately downstream of this sequence. Knowledge of nucleosome occupancy in the cell type of interest can also improve CRISPRi efficiency^53–55^ as accessibility of CRISPR/Cas9 complexes to the target site is key to success.

In a previous genome-wide CRISPR knockout study, we screened all protein coding genes in cattle to identify those that influence BHV-1 replication^49^. Having compiled a list of host genes with potential involvement in the viral replicative cycle, here we set about further evaluation of the 41 genes most likely to be involved in viral entry. To summarize, we validated the two receptors on the list, PVR and PVRL2, and their additive effect on mediating entry. In addition to genes coding for enzymes that directly catalyze biosynthesis of HS, we also confirmed the role the COG complex plays during virus entry, mostly likely by affecting cell surface levels of HS. In order to validate a broader range of candidate genes without undergoing the laborious process of generating and screening KO clones, we devised a multiplex CRISPRi system that enabled us to quickly test the candidacy of 27 other genes.

## Results

### Genome-wide CRISPRko screen identified many host genes important for BHV-1 entry into the cell

Our CRISPRko screen identified at least 41 pro-viral host genes related to cell entry. The proteins encoded by these candidate genes fall into three major categories: cell surface receptors, enzymes involved in GAG biosynthesis and proteins involved in trafficking within the Golgi apparatus (**Fig. 1, Fig. S1**). The two receptor encoding genes we recovered are LOC526865 (PVR or CD155) and PVRL2 (nectin-2) with the latter being suspected as a co-receptor but never experimentally confirmed ^24^. The screen also captured the prime importance of GAG for BHV-1 entry into the cell, with 22 of the candidates associated with Heparan Sulfate (HS) biosynthesis. In addition to enzymes known to catalyse HS synthesis^56^, other candidates have been found to affect HS levels and impact entry of viruses such as Chikungunya Virus^57^, Lassa virus^58^, Rift Valley Fever Virus^59^, and Vaccinia Virus^60^ (**Fig. 3A,** blue highlights). Some of these proteins are important for N-glycosylation, including Glycosyltransferases ALG5 and ALG10, and subunits of the Oligosaccharyltransferase complex OSTC and STT3A. Other candidates that may transport or catalyse formation of crucial precursor molecules for HS were also selected, namely GALE, NFS1, PTAR1, SLC35B2, SLC39A9^58,59^, UGDH and UGP2. In the third category, six genes encoding subunits of the COG complex^37^ (COG1, COG2, COG4, COG5, COG6, and COG7) were chosen. Other genes encoding resident proteins of the Golgi with demonstrated or suspected indirect roles in protein glycosylation^57–60^ (C1H3orf58, IMPAD1, MON2^61^, NAPG, SACM1L, TM9SF2, TM9SF3, TMED2, TMEM165 and UNC50) were also selected.

**Figure 1.**
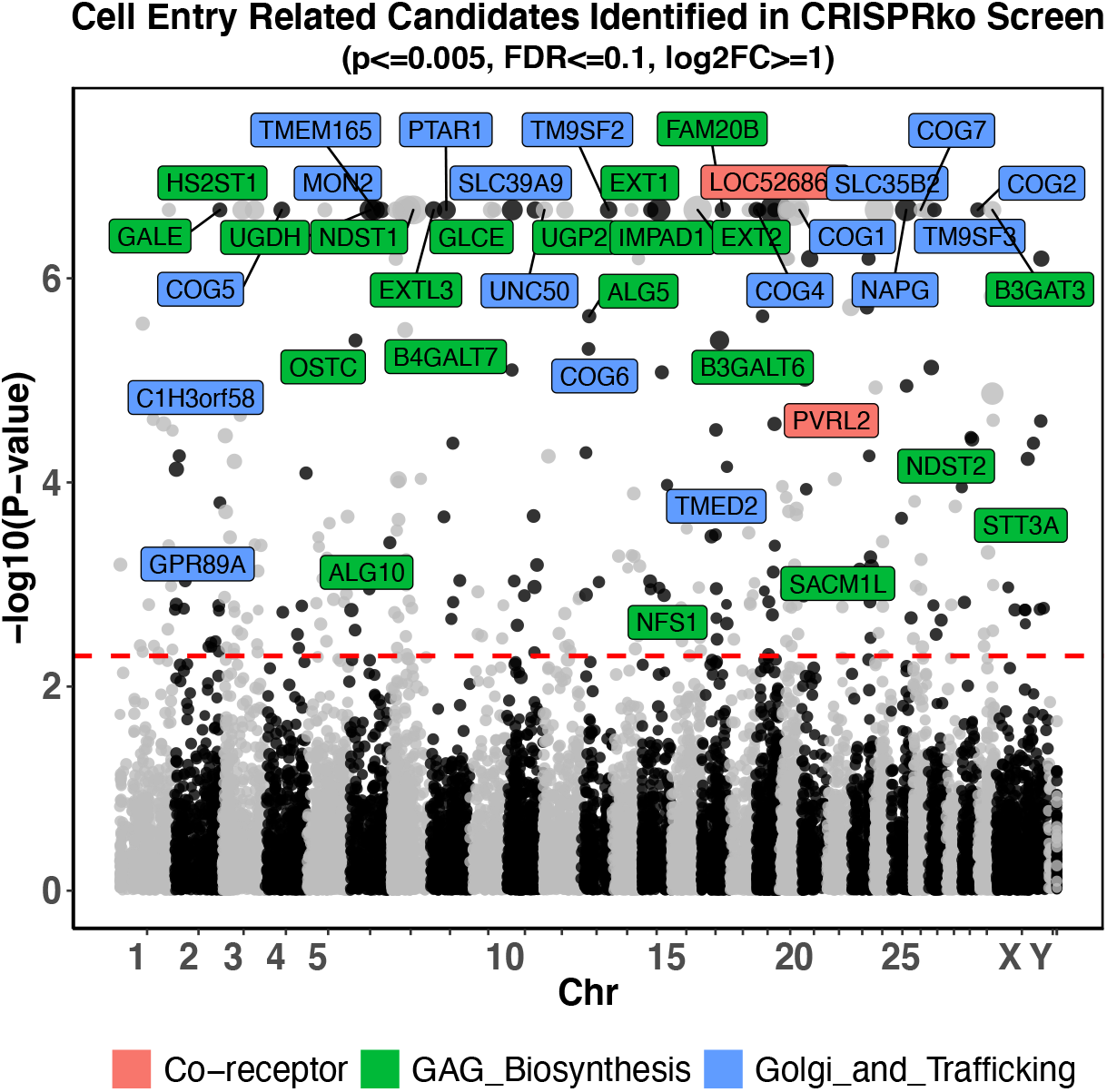
Genome-wide CRISPR knockout screen identifies pro-viral genes associated with viral entry into the cell. Guide RNA copy number changes obtained from the CRISPRko screen by comparing GFP Negative cells to GFP High cells^49^ were plotted. Each dot represents one of the 21,216 protein coding genes targeted by the btCRISPRko.v1 library, with its genomic location plotted against the x-axis and −log10(p-value) based on MAGeCK^79^ analysis against the y-axis; the sizes of dots represent −log2FoldChange values. Pro-viral candidate genes that are most likely to affect BHV-1 cell entry with p-value <= 0.005, False Discovery Rate(FDR) <= 0.1, and log2 fold changes (log2FC) >= 1 as statistical cutoff are highlighted; they are color coded based on their pathways to impact. The red line represents p-value threshold as 0.005 or −log10(p-value) = 2.3.

### Simultaneous knockout of PVR and PVRL2 severely impairs BHV-1’s ability to enter the host cell

While PVRL2 has been hypothesised as an entry receptor for BHV-1 its candidacy has never been properly tested. We first generated biallelic KO clones lacking this gene by transfecting *in vitro* transcribed sgRNA targeting PVRL2 into an MDBK cell line expressing Cas9 (Cas9+/+)^49^. By plaquing directly on these KO cells, we detected up to 1.9-fold decrease in virus titre and 2.7-fold reduction in plaque size compared to the parental Cas9+/+ cells (**Fig. 2A**). When we conducted the same experiment targeting PVR instead, the impact on the virus was less pronounced, with 1.7-fold and 2.1-fold drop in titre and plaque size respectively (**Fig. 2B**). Although the impact on virus replication caused by loss of either receptor alone was statistically significant, the degree of impact was rather modest. This is likely due to the ability of alpha herpesviruses to bind to multiple cell surface molecules to mediate entry. Next, we generated double KO (dKO) clones lacking expression of both PVRL2 and PVR and examined the impact on virus replication. Based on plaque assays, the impact of the dKO was more prominent than the sum of the individual gene KOs, with virus titre and plaque size reduced by up to 8.2-fold and 4.8-fold respectively compared to the parental Cas9+/+ controls (**Fig. 2C**).

**Figure 2.**
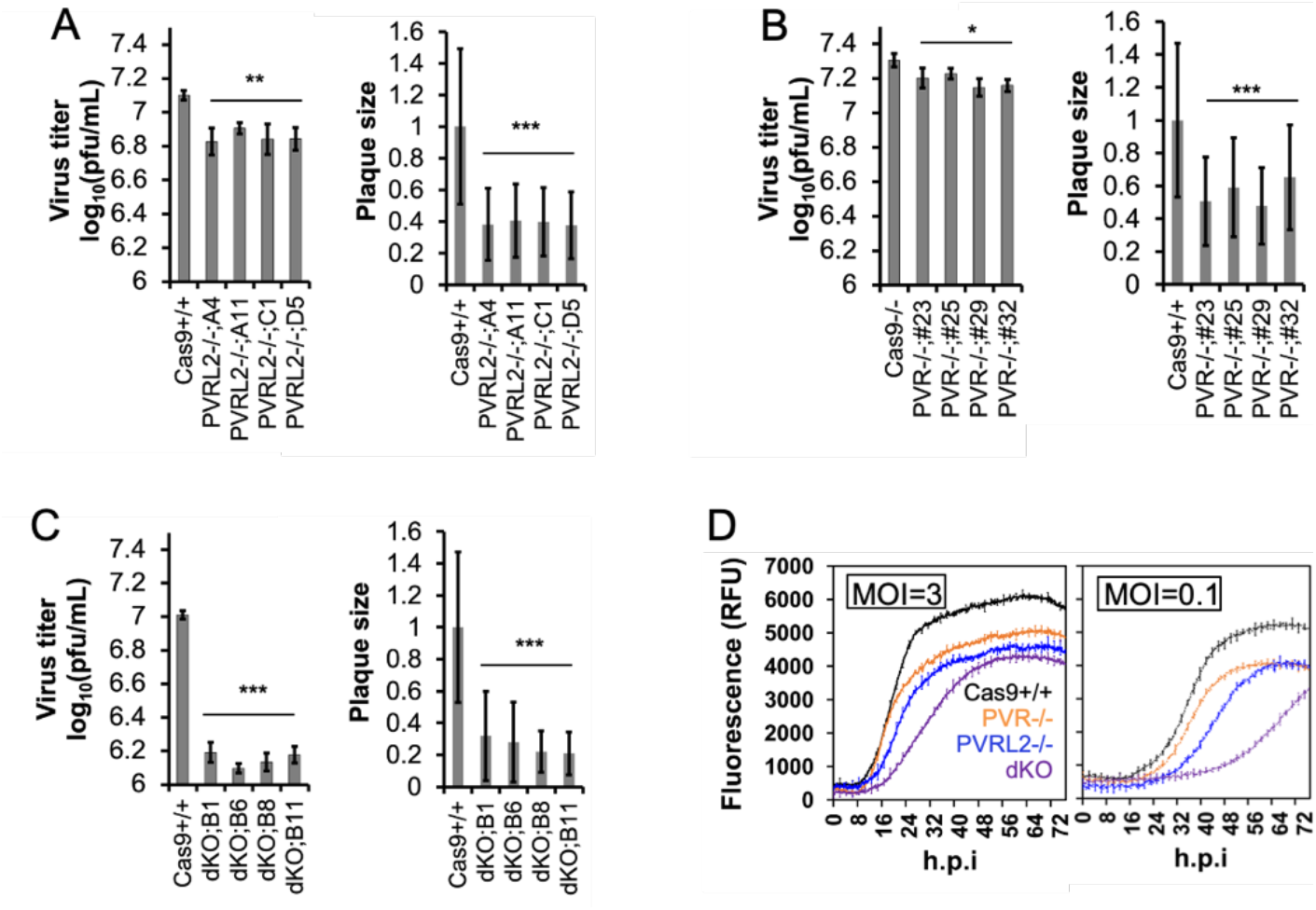
Loss of cell surface co-receptors impairs BHV-1 replication in MDBK cells. **A.** Plaque assay results in four clones with PVRL2 KO compared to Cas9+/+ controls cells. **B.** Plaque assay results in four clones with PVR (a.k.a LOC526865) KO compared to Cas9+/+ controls cells. **C.** Plaque assay results in four clones with PVRL2 and PVR double KO (dKO) compared to Cas9+/+ controls cells. For all plaque assays shown in **A**, **B**, and **C**: all plaque assays were done with at least two biological repeats (n >= 2) and plaque sizes from each condition were averages measured from at least 40 plaques and then normalized to that in Cas9+/+ cells; ***: p < 0.005; **: p < 0.01; *: p < 0.05 based two-tailed student’s t-test, error bars represent +/− 1 STD. **D**. VP26-GFP protein growth curve in Cas9+/+, PVR KO, PVRL2 KO, and PVR/PVRL2 double KO(dKO) infected with BHV-1 at MOI=3 or 0.1 measured as fluorescence (n = 4) throughout a 72-hour infection.

We then studied the rates of viral protein synthesis by sequential recording of GFP intensity in these receptor KO cells and controls for 72-hours post infection with a GFP tagged BHV-1^62^. We detected both a delay and a reduced magnitude of VP26-GFP expression in all KO cells (PVR−/−, PVRL2−/− and dKO) compared to the parental Cas9+/+, when cells were infected at either MOI = 3 or MOI = 0.1 (**Fig. 2D**). Taken together, these results confirm that both PVR and PVRL2 can serve as receptors for BHV-1 and they work cooperatively for the virus. Our data also consistently suggest that PVRL2 mediates more efficient entry than PVR in MDBK cells; guides targeting PVRL2 were more enriched in the CRISPRko screen^49^ and its loss lead to a more severer impact on viral replication manifested by greater loss of viral titre and spread (**Fig. 2A,B**) as well as bigger reduction and delay in viral protein synthesis (**Fig. 2D**).

### Heparan Sulfate is crucial for efficient BHV-1 entry

Like many other viruses, BHV-1 initiates entry by first interacting with cell surface HS. Out of the 41 viral entry associated candidates from the CRISPRko screen, at least 11 genes encode for enzymes that directly catalyse GAG linker assembly or HS chain initiation, elongation and modification. Four of these genes are crucial for steps of common chain linker assembly for both HS and CS/DS, include glycosaminoglycan xylosylkinase (FAM20B), galactosyl-transferases (B4GALT7 and B3GALT6), and Glucanosyltransferase (B3GAT3) (**Fig. 3A**). After linker assembly, the pathways begin to diverge, and different enzymes are required for HS and CS/DS chain initiation and elongation. The screen also identified EXTL3, one of the two Glycosyltransferases known to initiate HS chain formation and experimentally shown to be more efficient for this purpose than the other enzyme EXTL2 (**Fig. 3A**)^35,63^ . HS chain elongation is thought to be achieved by exostoses (EXT) genes EXT1 and EXT2, both candidate genes identified in our screen, with knockout of either of these genes completely abolishing HS synthesis^56^.

**Figure 3.**
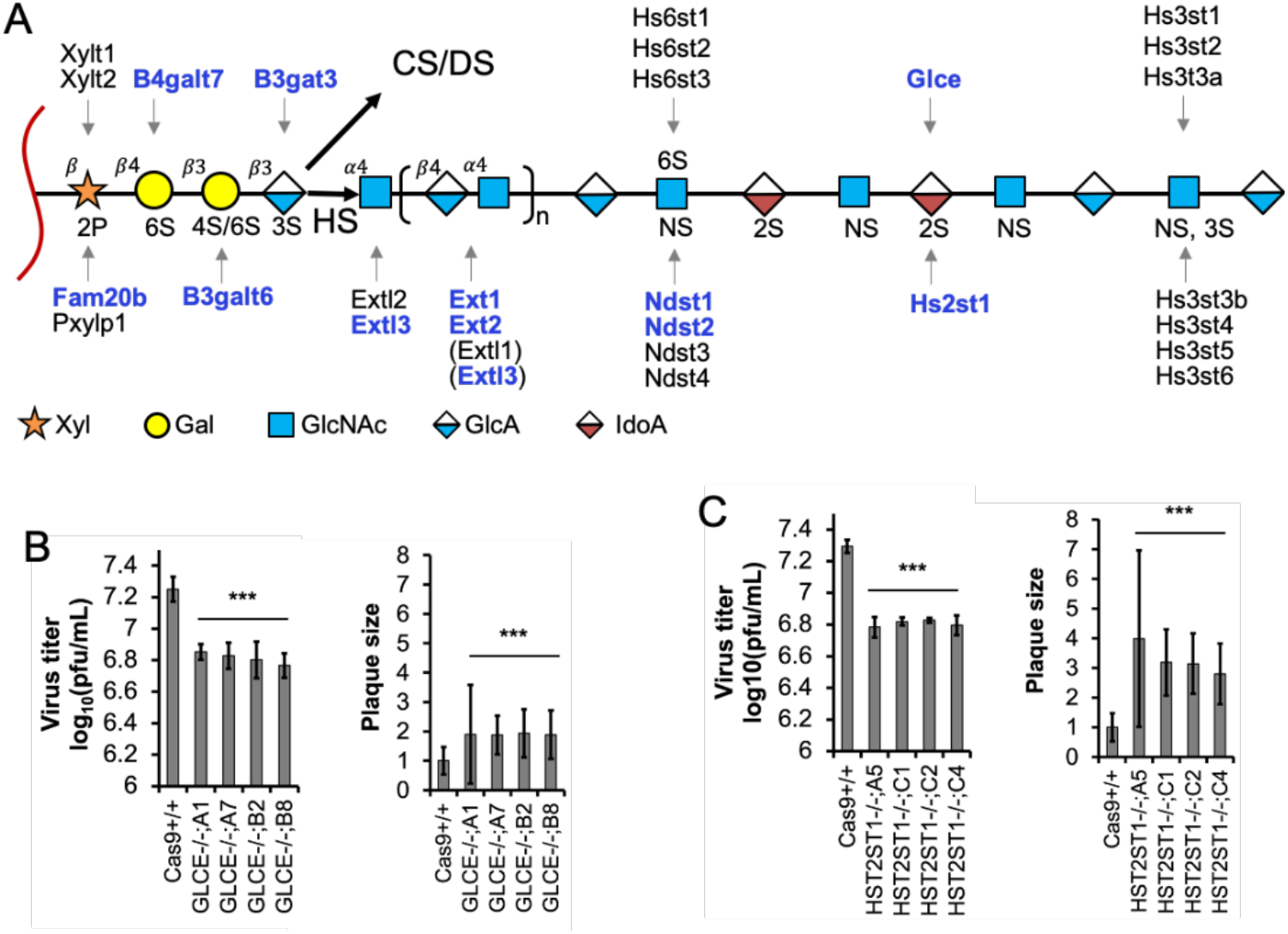
Many candidates catalyze cell surface Heparan Sulfate proteoglycan synthesis. **A.** Genes known to be involved in each step Heparan Sulfate (HS) biosynthesis and chain extension from the core protein (red squiggle); those highlighted in blue have been identified by the screen. This diagram is modeled after Figure 2 in Chen et al ^56^ with pathway to chondroitin sulfate (CS)/dermatan sulfate (DS) synthesis simplified. Abbreviations for units within the HS chain: Xyl, xylose residue; Gal, galactose residue; GlcNAc, 2-deoxy-2-acetamido-α-D-glucopyranosyl; GlcA, β-D-glucuronic acid; IdoA, α-L-iduronic acid. **B.** Plaque assay results from four GLCE knockout clones, A1,A7,B2 and B8 compared to Cas9+/+ control cells (n >= 6). Plaque size measurements were taken from at least 50 plaques for each clone and then normalized to Cas9+/+ cells. ***: p < 0.0001 based on un-paired t-test and error bars represent +/− 1 standard deviation (STD). **C.** Plaque assay results from four HST2ST1 knockout clones, A5,C1,C2 and C4 compared to Cas9+/+ control cells (n >= 5). Plaque size measurements were taken from at least 89 plaques for each clone and normalized to Cas9+/+ cells. ***: p < 0.0001 based on un-paired t-test, error bars represent +/− 1 STD.

As the HS chain polymerizes, the nascent heparosan chain undergoes a series of sulfation and epimerization carried out by four classes of sulfotransferases and an epimerase^25^. Our screen identified NDST1 and NDST2 (**Fig. 3A**), two of the four genes encoding homologous bifunctional N-deacetylase–N-sulfotransferases de-N-acetylates. While experiments in CHO cells showed that knockout of either gene alone did not lead to substantial reduction of disaccharide^56^ sulfation, when both genes were deleted simultaneously disaccharide sulfation was eliminated. In addition, the one epimerase that converts GlcA to IdoA (**Fig. 3A**) encoded by GLCE was also identified in the screen, along with HST2ST1, a 2-O-sulfotransferase that was predicted to interact with the protein product of GLCE^64^. To experimentally confirm the positive role of HS in cell entry, we first conducted Heparin blocking assays by treating cells with different concentrations of Heparin during a BHV-1 infection then titrating viruses produced from these cells on wild type MDBK cells; there was a steady decrease in virus titre as Heparin concentration increased (**Fig. S2**). We then produced KO clones lacking either GLCE or HST2ST1 and assessed viral replication in them. The loss of either gene alone led to average 2.7-fold (GLCE) and 3.1-fold (HST2ST1) reduction in virus titre based on plaque assay (**Fig. 3B, C**). Surprisingly, the plaque size was increased in both cases, up 1.9-fold in GLCE−/− cells and 3.3-fold in HST2ST1−/− cells compared to Cas9+/+ controls (**Fig. 3B, C**).

### COG knockout impairs virus replication

To evaluate the importance of the COG complex to virus replication, we first generated single gene KO clones of either COG6 or COG7. Plaque assays revealed that knockout of either gene reduced virus titre in multiple clones, with up to 4.3-fold and 1.5-fold decrease respectively (**Fig. 4A,B**). Interestingly, the sizes of plaques grown in COG6 −/− cells increased to at least 1.6-fold, whereas no significant augmentation was observed in COG7 −/− clones (**Fig. 4A,B**). A similar reduction in virus titre was also observed when the COG6 −/− cells were infected with Alcelaphine Virus 1, a gamma herpes virus endemic to wildebeest that causes malignant catarrhal fever when transmitted to cattle (**Fig. S3**)^65^. The negative impact on BHV-1 replication associated with the loss of COG6 expression was further evidenced by the delayed production of the VP26-GFP protein at high or low MOI compared to the Cas9+/+ control cells (**Fig. 4C**). To dissect this phenomenon, we harvested supernatant and cell pellets from the virus infected COG6 −/− cells and titrated these on wild type MDBK cells. There were fewer infectious viral particles in both the supernatant and cell pellets collected from infected KO cells compared to parental Cas9+/+ cells by 24 h.p.i (**Fig. 4D,E**), with the difference becoming greater when infection was extended to 72 h.p.i (**Fig. 4E**).

**Figure 4.**
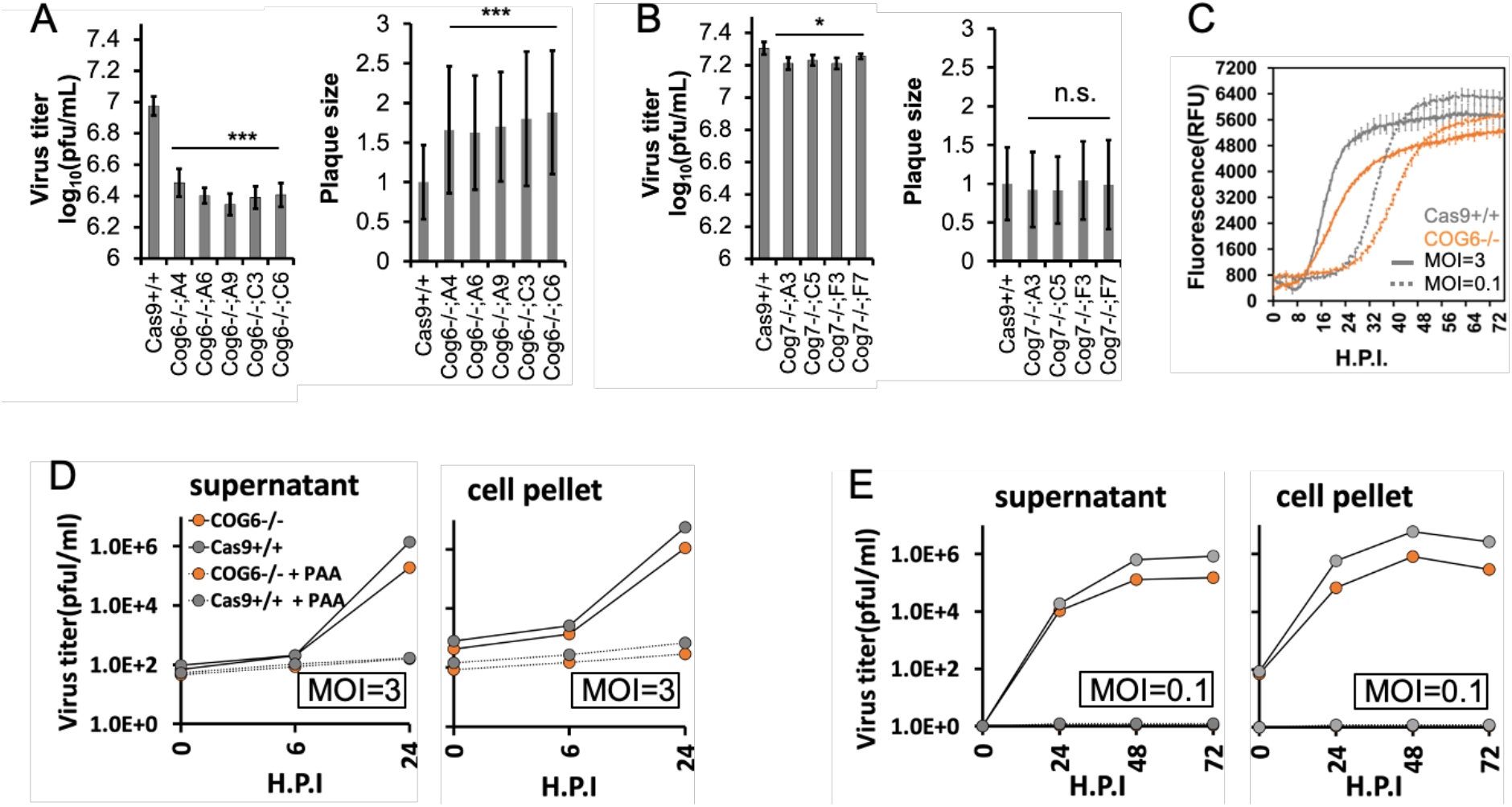
The loss of COG6 and COG7 affects BHV-1 replication in MDBK cells. **A.** Plaque assay results from five COG6 KO clones compared to Cas9+/+ control cells. B. Plaque assay results from four COG7 KO clones compared to Cas9+/+ control cells. For both **A** and **B**, viral titers were measurements from at least three repeats (n >= 3); plaque sizes were averaged from >= 65 plaques for each condition and then normalized to that from Cas9+/+ cells. ***: p < 0.005; *: p < 0.05; n.s.: not significant based on unpaired two-tailed student’s t-test, error bars represent +/− 1 STD. **C.** VP26-GFP fluorescence growth curve in Cas9+/+ and COG KO cells infected with GFP tagged BoHV-1 at MOI=3 or 0.1. **D.** Plaque assay results by tittering virus harvested from COG6 KO or Cas9+/+ cells infected with BoHV-1 at MOI=3 on wt MDBK cells, supernatant and cell pellet samples were collected at 0,6,24 h.p.i. PAA was added 1 hour prior to infection to serve as negative control. Viral samples were collected from a single infection and the titration was repeated twice (n = 3), the error bars are too small to be shown. **E.** Experiment conducted as in **D** but with MOI=0.1 and samples were harvested at 0,24,48 and 72 h.p.i.

### COG6 knockout reduces abundance of viral genomes and viral transcripts

To help determine the BHV-1 replicative step most affected by COG6 KO, we surveyed the abundance of viral nucleic acid in COG6 −/− cells. We first quantified total viral genomic DNA(gDNA) in cells six hours after infections using qPCR. Viral genome copy numbers from COG6 −/− cell pellets were 2.2-fold (MOI = 3) and 3.8-fold (MOI = 0.1) lower compared to either the parental Cas9+/+ or wt MDBKs (**Fig. 5A**). Since BHV-1 completes one cycle of replication in 6-8 hours, the decline of viral gDNA copies in these KO cells suggests impairment to either entry, capsid trafficking to the nucleus or genome replication within the nucleus. To narrow down these possibilities, we next assayed levels of viral transcripts from all three stages of viral transcription in these infected cells, immediate early (IE), early (E), and late (L). Transcripts of ICP4, an IE gene encoding a master transcription factor for many viral genes, were lower in the COG6 −/− cells by 2-fold (**Fig. 5B**). As IE genes are transcribed in the nucleus directly from the original viral genome, prior to de novo viral gDNA synthesis, reduction in ICP4 transcripts indicates an impediment to viral genomes reaching the nucleus and therefore either comprimised entry or retro-grade trafficking, assuming transcription efficiency of IE genes is not affected by COG6 KO. Transcripts from an E gene encoding thymidine kinase, UL23, were also lower in COG6 −/− cells by 4-fold (MOI =3) and 2.5-fold (MOI =0.3) (**Fig. 5C**), while the greatest impact was observed in the L gene transcripts of VP26 whichh were supressed by 14-fold (MOI =3) and 7.1-fold (MOI = 0.3) in COG6 −/− cells compared to the parental Cas9+/+ (**Fig. 5D**). Taken together, the data support loss of COG6 expression impacting the efficiency with which BHV-1 genome reach the nucleus, with knock-out effects on both transcription and genome replication.

**Figure 5.**
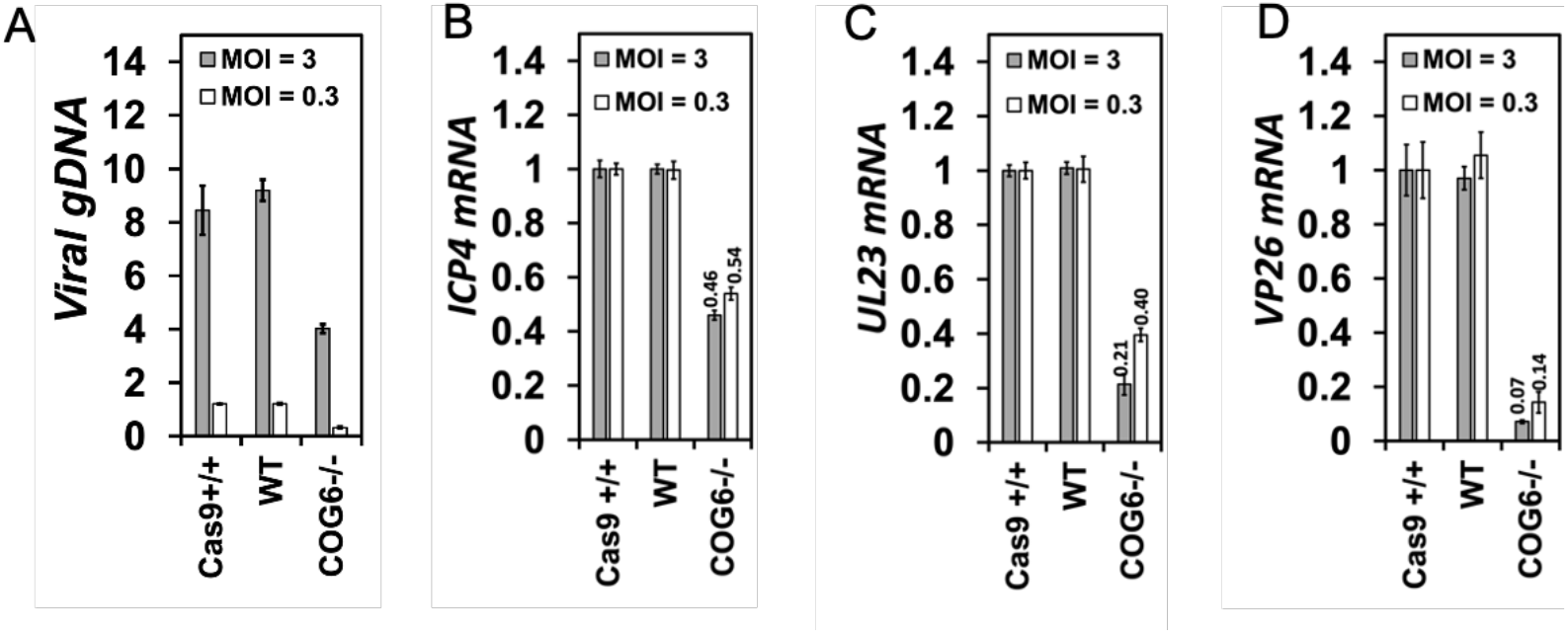
COG6 KO leads to reduction of viral genomic DNA and viral transcript abundance in MDBK cells. **A.** Total relative viral genomic DNA(gDNA) in Cas9+/+, WT and COG6 KO cells at 6 h.p.i at MOI=3 or 0.1, measured by qPCR with the levels in Cas9+/+ cells infected at MOI of 0.1 set as 1. **B.** Relative ICP4 mRNA levels in Cas9+/+, WT, or COG6 KO cells infected with BHV-1 at MOI=3 or 0.1 at 6 h.p.i measured by qPCR following reverse transcription. **C.** Experiment conducted as in B but UL23 mRNA level was measured instead. **D.** Experiment conducted as in B but VP26 mRNA level was measured. **Note**: all data shown here were obtained from a single infection experiment with three q-PCR technical repeats.

### CRISPRi leads to efficient knockdown of host gene transcription

The optimized protocol we devised to produce single cell KO clones for the validation of candidate genes is an extremely effective process but remains time consuming and labour intensive. From initial *in vitro* transcription of gene specific sgRNA to the analysis of Sanger sequencing to genotype expanded clones takes approximately one month. For single genes of specific interest this is a cost-effective process but is not compatible with biological assessment of larger candidate gene lists. Additionally, complete ablation of gene function is not feasible for candidate genes that are also essential for cellular fitness, often manifested by failure to obtain KO clones. To overcome these limitations, we produced an alternative cell line designed for CRISPRi induced gene knockdown (KD) (**Fig. 6A**) by modifying the rosa26 locus with a dCas9-KRAB-MeCP2 fusion protein expressed from a doxycycline inducible TetOn promoter (**Fig. 6B, C**). Using the same targeting and genotyping strategies employed to make the Cas9 expressing cell lines for the screen (**Fig. 6C**)^49^, we created a homozygous dCas9+/+ cell line for CRISPRi which was confirmed by genotyping PCR (**Fig. 6D**, second lane).

**Figure 6.**
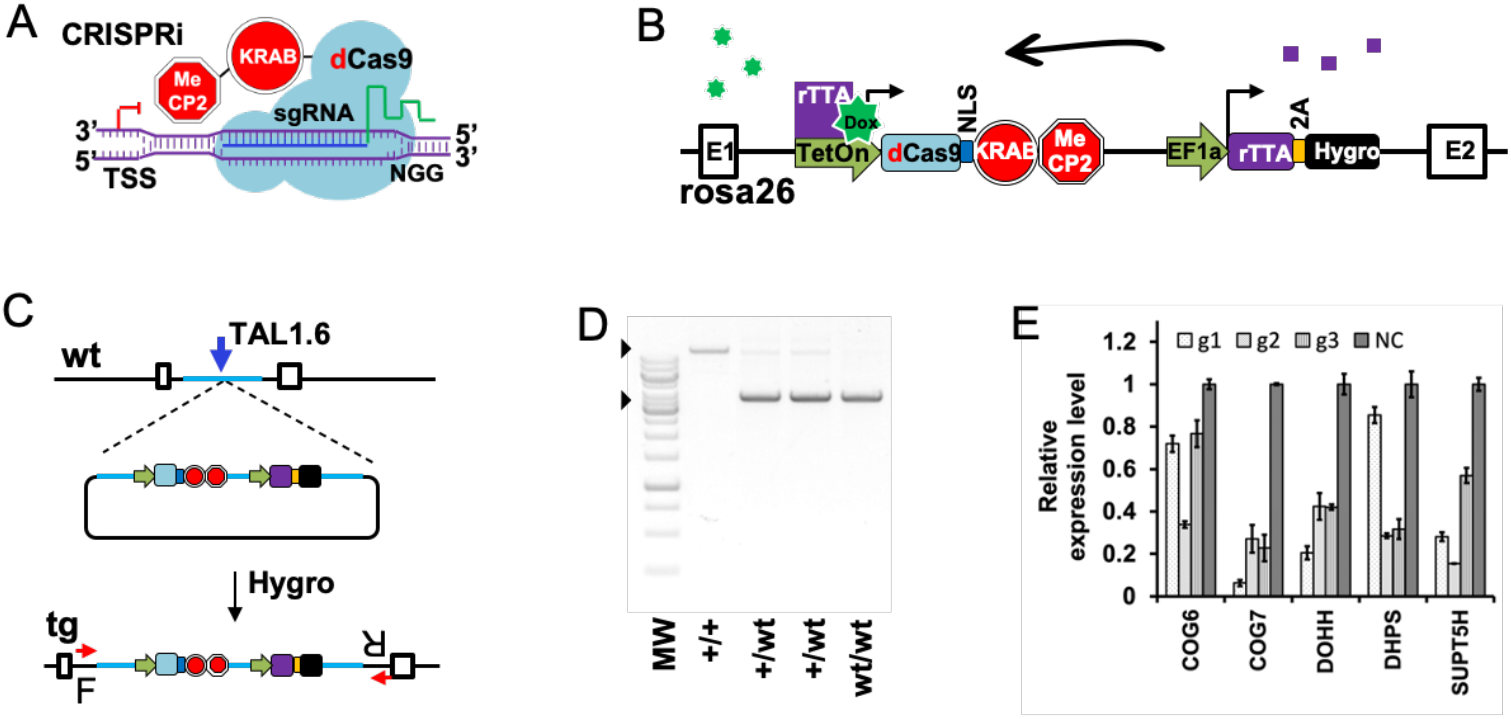
CRISPRi mediates efficient knockdown of gene expression in MDBK cells. **A.** The CRISPRi system applied in this study, based on the module developed by Andrea Califano et al (See **Materials and Methods**). **B.** Doxycycline inducible expression of the dCas9-KRAB-MeCP2 module from a stably integrated expression cassette at the cow rosa26 locus. Green heptagram represents Doxycycline, purple squares are rTTA expressed from the downstream expression cassette and is used to enhance Doxycycline induction. **C.** Targeting strategy to knock-in the dCas9-KRAB-MeCP2 cassette into rosa26 using TAL1.6 stimulated HDR)^49^ and Hygromycin selection, prior to dilutional cloning and genotyping by PCR using primer set F+R (red arrows). wt: wild type; tg: targeted. **D.** Representative genotyping results by PCR from a homozygous targeted clone (+/+), and two heterozygotes (+/wt), black arrows point to amplicons from the targeted allele (top) and wt allele (bottom). **E.** Relative expression levels of five genes in dCas9+/+ MDBKs targeted by CRISPRi (g1,g2,g3) compared to cells transfected with non-targeting CRISPRs (NC) based on RT-qPCR results from a single experiment with three technical qPCR repeats, error bars represent +/− 1 STD.

Using these cells, we initially targeted five candidate genes, COG6, COG7, DOHH, DHPS, and SUPT5H, by introducing single guides and measuring transcript levels by reverse transcription and qPCR. Three guides, designed to bind within a 400bp region immediately downstream of the TSS of each gene, were cloned individually into PiggyBac based sgRNA expression vector PB-U6g5_PGK_Puro2aBFP (**Materials and Methods**). This vector was then co-transfected into dCas9 +/+ cells with the transposase plasmid pCMV-hypBase^66^, allowing efficient stable transgene integration by transposition. Following puromycin selection for stable integrants, cells were cultured with 500ng/ml doxycycline for 48 hours prior to mRNA isolation. Compared to control cells transfected with a non-targeting guide, candidate gene expression levels fell to 6-85% of the basal levels in cells containing one of the three targeting guides (**Fig. 6E**). For each gene at least one of the guides resulted in transcript suppression to below 40% of basal level. These results demonstrate that this CRISPRi system is very effective in suppressing gene expression in most cases.

### CRISPR3i enables rapid validation of candidate genes

We reasoned that simultaneous expression of three guides tiled along the 400bp downstream of a TSS (**Fig. 7A**) would have a synergistic impact on candidate gene transcript suppression. In order to achieve reliable, coordinated expression of all guides in a transfected cell pool they must all be delivered using a single construct. We therefore devised a one-step cloning protocol (**Supplementary file 1**) that quickly assembles three sgRNA expression cassettes into the same PiggyBac based sgRNA expression vector used previously (**Fig. 7B**). For the ligation, the linearized PiggyBac vector was mixed with three CRISPR containing short dsDNA fragments assembled from annealed oligonucleotides and two pre-cut fragments supplying sgRNA scaffolds and small RNA promoters hH1 and mU6^67,68^. Unique overhangs at either end of fragments ensured orientation specific assembly. The resulting CRISPR3i vectors harbour an array of three sequential sgRNA expression cassettes, each with a unique promoter.

**Figure 7.**
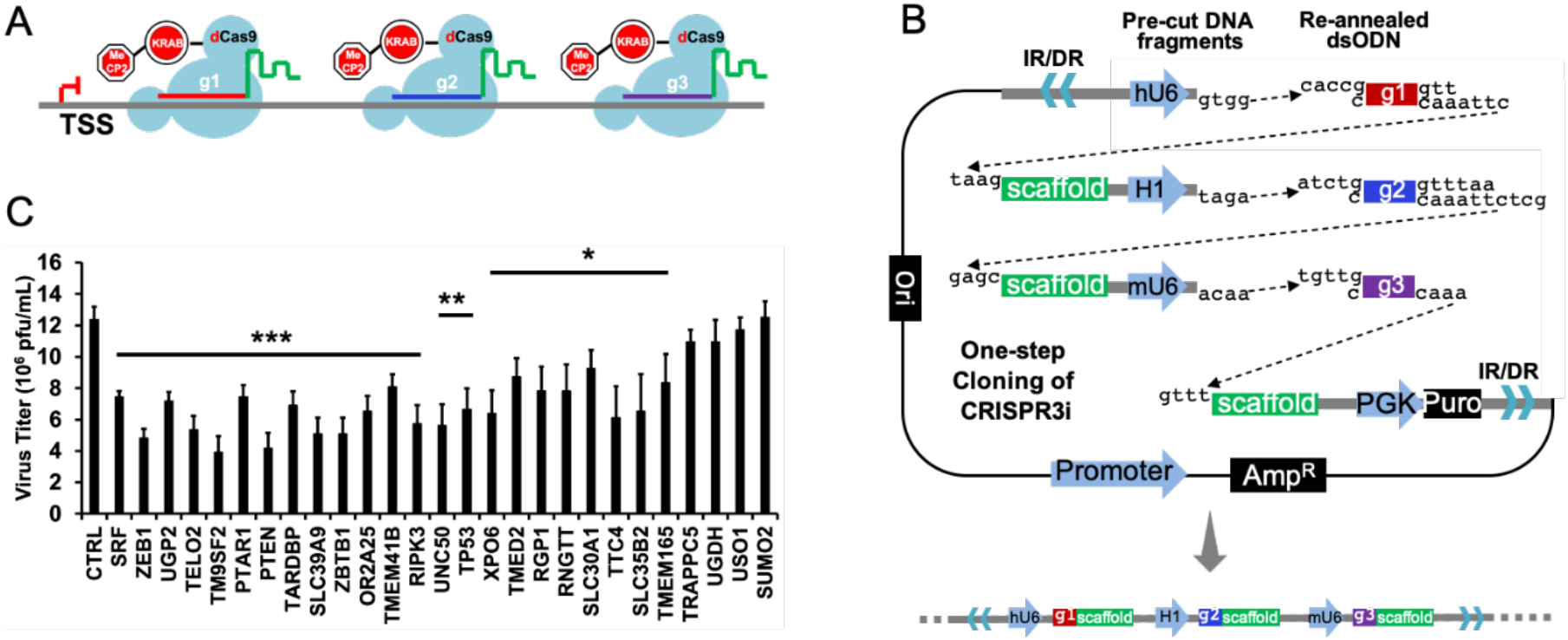
Multiplex CRISPRi against pro-viral candidate genes results in reduced viral titers. **A.** The one-step cloning strategy to synthesize PiggyBac transposon vectors carrying three sgRNA expression cassettes in tandem to implement CRISPR3i by PB transposition. **B.** CRISPR3i against a host gene in action by binding immediately downstream of TSS in tandem for synergistic gene repression. **C.** Plaque assay results in cells with CRISPR3i expression intended to interfere with transcription of specified pro-viral genes involved in HS biosynthesis and other functions identified by the CRISPRko screen; CTRL represents virus titer from cells expressing three non-targeting sgRNAs (n >= 3). Results were ordered by p-value. ***: p < 0.005; **: p < 0.01; *: p < 0.05 based on two-tailed student’s t-test, error bars represent +/− 1 STD.

We used CRISPR3i to assess 27 candidate genes, including ten associated with GAG biosynthesis. We constructed 28 CRISPR3i PiggyBac vectors, one targeting each candidate gene plus a negative control vector. CRISPR3i constructs were each co-transfected with pCMV-hypBase into dCas9+/+ MDBK. Cell pools were selected with puromycin then induced with doxycycline 48 hours prior to plaque assay, with the addition of doxycycline to the overlay throughout the incubation period to maintain consistent host gene suppression. Analysis revealed that the CRISPR3i approach resulted in significant reduction of virus titre in 23 out of the 27 candidates (**Fig. 7C**). The protein products of these genes directly or indirectly affect HS biosynthesis (**Fig. 1**), regulate cell cycle progression (PTEN, SRF, TELO2), apoptosis (RIPK3), transcription and RNA processing (RNGTT, TARDBP, ZBTB1), with additional genes having as yet uncategorized roles (OR2A25, RGP1, SLC30A1, TMEM41B, TTC4, XPO6, ZEB1) (**Fig. 7C**). These results confirm the pro-viral roles of most of the genes tested and demonstrates the utility of CRISPR3i in studying host pathogen interactions by modulating host gene expression.

## Discussion

In this study, we experimentally validated many candidate genes related to viral entry identified by our previous genome wide CRISPR KO screen. For the first time, we prove that PVRL2 is indeed a receptor for BHV-1 entry into bovine cells. We also found that it likely mediates more efficient entry than the previously identified PVR, as PVRL2 KO had a bigger impact on virus replication than did PVR KO (**Fig. 2**). While double KO of both PVR and PVRL2 further reduced viral replication, it failed to confer MDBK cells with complete resistance to infection. Previous studies demonstrated that by editing a single cell surface receptor, pigs could be engineered to become resistant to porcine reproductive and respiratory syndrome virus^69,70^ and transmissible gastroenteritis virus^71^ infection, but our data show that this is not an appropriate strategy to create animals that are fully resistant to BHV-1. This was likely due to the presence of other surface co-receptors such as nectin-1 for BHV-1 and/or entry via receptor independent mechanisms.

Our CRISPRko screen identified key enzymes involved in multiple steps of HS synthesis (**Fig 3A**), and here we confirmed the significance of cell surface HS in mediating BHV-1 entry. Interestingly, MDBK clones missing HS2ST1 or GLCE produced fewer plaques as expected, but surprisingly those that were formed were bigger compared to the parental controls. One possible explanation is that while lack of HS results in fewer initial infections and thereby lower titres, once an infection becomes established newly synthesized viral particles distribute more widely in the plate due to the reduced electrostatic attraction to the plasma membrane of adjacent cells. Indeed, it has been observed that multiple herpes viruses including HSV-1, HSV-2 and BHV-1 upregulate heparinase transcription and expression, thereby promoting viral release^72,73^. The 3-O-sulfated disaccharides in HS are created by seven (**Fig. 3A**) glucosaminyl 3-O-sulfotransferases in mammals, six of which are known to generate a binding site for gD of HSV-1 in humans^30^. This high functional redundancy is the likely reason why none of these genes was identified by the KO screen. In addition, despite identifying many genes crucial for HS production, no enzymes specific for CS chain initiation, extension or modification showed up in the screen^56^. This indicates that even though CS has been shown to play an auxiliary role in cell surface binding by HSV-1 ^32,74^, it is not required for BHV-1 to mediate entry into MDBK cells, a conclusion supported by our chondroitin sulfate blocking assay (**Fig. S2**).

Another prominent cluster of candidates revealed by the screen encode six subunits of the COG complex. Reduced viral titre and delayed viral replication in COG6 −/− and COG7 −/− MDBK cells verified its pro-viral roles in BHV-1 replication (**Fig. 4**). We also detected a decrease in total viral gDNA and transcripts in COG6 KO cells at 6 h.p.i after a single cycle of replication (**Fig. 5**), suggesting impairment to either host cell entry or capsid trafficking to the nucleus. Interestingly, as with HST1ST2 and GLCE KO cells, plaques associated with COG6 KO cells were bigger than in the parental controls; the reduced cell entry but faster release of BHV-1 in these KO cells strongly indicates disrupted HSPG biosynthesis. In addition, we observed a more pronounced reduction in viral titre following COG6 KO versus COG7 KO (4.3-fold vs 1.5-fold respectively), and a more stressed cellular morphology in the former (data not shown). The COG complex is formed by two lobes, with lobe A containing subunits COG1-4 and lobe B composed of COG5-8^36^. Each subunit has been found to interact with overlapping yet non-identical subsets of Rab GTPases, golgins, and SNAP receptors (SNAREs)^36^, implicating differential outcomes from individual subunit KO. As the COG complex participates in multiple aspects of the Golgi functionality such as intra-cellular trafficking and protein glycosylation, our data do not rule out the possibility that in addition to HS deficiency, COG complex perturbation could result in additional cellular defects that contribute to impaired virus replication. By comparing GLCE KO cells with COG deficient cells, one study observed similar profiles of glycosylation impairment in the two lines but also glycosylation-independent cellular defects in the latter, including fragmented Golgi, abnormal endolysosomes, defective sorting or delayed retrograde trafficking^75^.

Finally, we demonstrated that CRISPRi can be a powerful tool when applied to modulate host gene expression in the context of a viral infection. To mitigate gaps in our knowledge relating to accurate TSS and nucleosome occupancy information for cattle, we developed an easy to assemble multiplex CRISPRi system, CRISPR3i, that utilises three guides simultaneously. Using this approach, we successfully validated 23 out of 27 candidates obtained from the CRISPRko screen. The failure to validate the remaining four candidates does not necessarily indicate false positives as the KD threshold required for phenotypic consequence may not have been reached. Here we used CRISPR3i to target a single gene at a time, but an alternative approach could be to employ guides designed to different genes but expressed from a single vector. It would be relatively simple to further expand this system to deliver a greater number of guides simultaneously.

The full potential CRISPRi can offer to large animal research remains to be explored. RNA interference (RNAi) primarily degrades or blocks the translation of transcripts that are exported to the cytoplasm, leaving those that reside and function in the nucleus unaffected. This means that many viral or host non-coding RNAs cannot be adequately interrogated by RNAi. CRISPRi by contrast can be designed to repress the expression of almost any gene, regardless of the destination of the transcripts. CRISPRi is also an excellent alternative to KO when targeting genes essential for cell fitness or those from large gene families with high sequence similarity between members. Furthermore, the inducible nature of CRISPRi allows the level of KD to be tuned by varying the doxycycline dose (**Fig. S4**). In the current study we utilised CAGE data from water buffalo^76^ to refine our bovine TSS prediction, although less than 30% of these CAGE tags mapped to the vicinity of annotated promoters on a genome wide level (unpublished data). Ongoing efforts to better annotate a plethora of genomes will inevitably contribute to improved CRISPRi design.

## Materials and Methods

All procedures or materials, if not listed here, such as nucleic acid extraction, RT-qPCR, and VP26-GFP growth monitoring with CLARIOStar, are available in the **Supplement** or described in detail previously^49^.

### Cells and viruses

Wild type Madin-Darby bovine kidney cells (MDBK), Cas9+/+ MDBK cells or dCas9+/+ cells49 were cultured in DMEM (D5796, Sigma) supplemented with 5% horse serum (26050088, Gibco), 1% Pen/Strep (15140122, Gibco), 1% L-glutamine (25030024, Gibco), 1% NEAA (11140035, Gibco), 1% sodium pyruvate (11360039, Gibco). Cells were kept in an incubator set at 37 °C with 5% CO2. A GFP tagged BHV-1 strain based on strain Jura^62^ was used throughout the experiment.

### Plasmids

The dCas9 targeting vector was created by Gibson Assembly, incorporating the inducible dCas9-KRAB-MeCP2 fusion cassette from PB-TRE-dCas9-KRAB-MeCP2 (gift from Andrea Califano, Addgene # 122267), the EF1a promoter driven rtTA fragment from PB-CA-rtTA Adv (gift from Andras Nagy, Addgene # 20910), and an in-frame Hygromycin selection marker connected by 2A peptide sequence. These cassettes were flanked by ~1kb left and right homology sequences amplified from the bovine rosa26 locus to enable HDR mediated knock-in. The PiggyBac vector used to deliver CRISPRi, PB-U6g5_PGK_Puro2aBFP, was constructed by cutting out the segment containing the hU6 sgRNA expression cassette and the Puro2aBFP selection marker from pKLV2-U6gRNA5(BbsI)-PGKpuro2ABFP-W (gift from Kosuke Yusa, Addgene # 67974) and cloning it into a PiggyBac backbone. The transposase expression vector pCMV-hypBase was shared by Dr. Kosuke Yusa while he was based at the Wellcome Sanger Institute. The H1 and mU6 donor plasmids were created by inserting PCR fragments with scaffold and H1/mU6 sequences into the Zero Blunt Topo vector.

### dCas9 knock-in to the rosa26 locus

The dCas9 fusion protein expression cell line dCas9+/+ was created by transfecting cells with 10ug of the targeting vector and 1ug TALEN 1.6 mRNA described previously^49^. After 3 days of culture at 33°C, cells were selected with 500ng/ml hygromycin. The process of isolating and expanding targeted single cell clones was completed as described previously^49^.

### Guide RNA design

Guides used for generating the KO clones were selected from the btCRISPRko.v1 library^49^. The CRISPRi/3i guides were designed using a similar pipeline to that used for the btCRISPRko.v1, utilising the 400bp genomic sequence immediately downstream of the TSS rather than protein coding sequence. The TSS information were extracted and combined from both the Ensembl (release-95) and NCBI (GCF_002263795.1) annotations of the Bos taurus assembly ARS-UCD1.2. Adjustments were made by mapping CAGE sequencing data from water buffalo onto the bovine genome using CAGEr^76,77^. For genes with two TSS annotations within 1 kb of each other, the downstream TSS was chosen. For genes with the two predicted TSS annotations greater that 1 kb apart, the NCBI annotation was used for the design. Please refer to **supplementary Table S1** and **supplementary file 2** for guide RNA sequences.

### CRISPR3i vector cloning

To prepare the fragments, the dsODNs containing the guides were annealed by mixing two complementary oligonucleotides and cooling from 95°C to 25°C at 0.1°C/second; the vector backbone and H1 mU6 donor plasmid were digested with BbsI and fragments of the predicted sizes were gel extracted. The six fragments (**Fig. 4**) were ligated using T4 ligase then transformed into Stabl3 competent cells. Colonies were pre-screened with bacterial PCR and Sanger sequencing verified before plasmid purification. For a more detailed protocol for the cloning, please refer to **supplementary file 1**.

### Transfections

All transfections were done using a Neon electroporator with parameters as follows: voltage, 1200v, duration: 30ms, pulses, 2. To generate KO clones, 5 × 10^5^ Cas9+/+ cells were loaded into a 100ul Neon tip and electroporated with 1ug of gene specific sgRNA transcribed *in vitro* as previously described^49^; for CRISPRi/3i targeting, PB vector carrying CRISPRi expression cassettes and pCMV-hypBase were mixed at 3ug:1ug ratio prior to electroporation.

### RT-qPCR

Sample extraction, reverse transcription and qPCR were conducted as before, using primers listed in **Table S2**.

### Generation of KO clones and stable cells with CRISPRi/3i knockdown

Cas9+/+ cells was transfected with 1ug *in vitro* transcribed guide and after three days of culture, cells were plated at 50-100 cells/10cm dish. A week after plating, clones were picked, expanded and genotyped by Sanger sequencing and TIDE analysis^78^ (**Table S3**).

### Viral infections and plaque assays

Briefly, viral stocks were defrosted in a 37°C water bath and diluted in full growth media with 2% horse serum. 1ml of diluted virus was added to each well of a 6-well plate and rocked gently to evenly distribute the virus. 1 hour after the addition of virus, the viral supernatant was removed and 2ml of fresh medium supplemented with 5% horse serum was added; this time point was regarded as zero hours post infection or 0 h.p.i. For plaque assays, 2mls of medium with 2% horse serum and 0.5% Avicel was added at 0h.p.i instead and the plates were incubated for four days prior to fixation and staining. The plates were scanned using an Epson document scanner and the sizes of plaques measured using ImageJ software. To assess titre of virus grown in infected cells, supernatant and cell pellet samples were harvested from each well at specified time points. The supernatant was used to infect non-modified Cas9+/+ cells directly, whereas the pellets were resuspended in 500ul PBS, frozen and thawed once and the spun supernatant used for plaquing.

### Statistical analysis

Pairwise comparisons were done using two-sided Student’s t-test.

## Acknowledgements

This study is funded by BBSRC grant BB/P003966/1, and BBSRC strategic funding to the Roslin Institute via grants BB/P013740/1 and BB/P013759/1. The authors would like to thank Drs. Andrea Califano, Andra Nagy and Kosuke Yusa for plasmids shared via Addgene as well as Dr. Peter Wild and colleagues from ETH for the GFP tagged BHV-1 virus. We are grateful to Professor David Hume and members from his lab, Rachel Young and Lucas Lefevre, for sharing their water buffalo CAGE data. We are also very appreciative of the advice and guidance given by Dr. Lel Eory, also based at the Roslin Institute, on the CAGE data analysis.

## Author contributions

**W.S.T:** Funding, Conception of studies, Methodology, Formal data analysis, Writing-draft, review and editing; **E.R.:** Methodology, Data analysis, Writing-review and editing; **I.D.:** Methodology, Data analysis, Writing-review and editing; **S.L.**: Funding, Writing-review and editing**; A.L.**: Funding, Writing-review and editing; **B.W.:** Funding, Writing-review and editing**; R.G.D.:**Funding, Conception of studies, Data analysis, Writing-review and editing.

