## supplementary file 1 for "Host cell factors important for BHV-1 cell entry revealed by genome-wide CRISPR knockout screen"

1. Digest 10ug of pKLV2-U6gRNA5(BbsI)-PGKpuro2ABFP-W (referred to as PB_3i from now on, Addgene #67974) >=4hrs, gel purify vector and elute with 50ul H2O to obtain linear vector. Alternatively, use uncut PB_3i in Step3 directly but background may be higher if digestion is not complete.
2. Assemble the re-annealing reactions below;

| Components | Volume (uL) |
| --- | --- |
| sgRNA sense + anti-sense oligo (50 µM) | 1 |
| 10X NEB Buffer 2 | 0.5 |
| ddH2O | 3.5 |
| Total | 5 |

a. Anneal the oligos in a thermocycler by using the following parameters: 95 °C for 5 min; ramp down to 25 °C at 0.1°C/s, 4°C hold;
b. Mix 1ul of re-annealed oligos from each g1, g2 and g3 with 97ul nuclease free H20 for a 1:100 dilution;
c. Proceed with BbsI digest and ligation;
3. Digestions and ligation:
d. Assemble the reactions in strip tubes below, replace oligo mix with H2O as negative control:

| Components | Volume (uL) |
| --- | --- |
| PB_3i (~50 ng/ul, linear or circular) | 0.25 |
| TOPO_g1tracr_H1(100ng/ul) | 0.5 |
| TOPO_g2tracr_mU6(100ng/ul) | 0.5 |
| Diluted oligo duplex mixes from above | 1 |
| 10X CutSmart buffer | 1 |
| NEB BbsI(10u/ul) | 0.5 |
| Quick T4 ligase(2000u/ul) | 0.25 |
| 10mM ATP | 0.5 |
| 20mM DTT | 0.25 |
| ddH2O | 5.25 |
| Total | 10 |

e. Incubate reactions in a thermo cycler running the following program: 10 X (37**°**C, 5 min; 16**°**C, 10 min); 37**°**C, 30 min + 65**°**C, 10 min.
4. Assemble the following plasmid-safe nuclease treatment and incubate reaction at 37**°**C for 30 min:

| Components | Volume (uL) |
| --- | --- |
| Reaction from above | 10 |
| PlasmidSafe buffer | 1.5 |
| H2O | 1.75 |
| 10mM ATP | 1.25 |
| Plasmid-Safe nuclease | 0.5 |
| Total | 15 |

5. Transform 2ul of the above reactions into 25 ul of Stbl3 or NEB Stable competent cells and plate the transformed E. Coli onto Carb^50^ or Amp^100^ plates.
6. The second day, pick colonies for bacterial PCR and start 5ml of culture for minipreps using Qiagen miniprep columns and Plasmid plus buffers with endotoxin removal. Elute with 30ul of endotoxin-free, EB, TE or H2O.
