## supplement for "Host cell factors important for BHV-1 cell entry revealed by genome-wide CRISPR knockout screen"

**Supplementary Data**

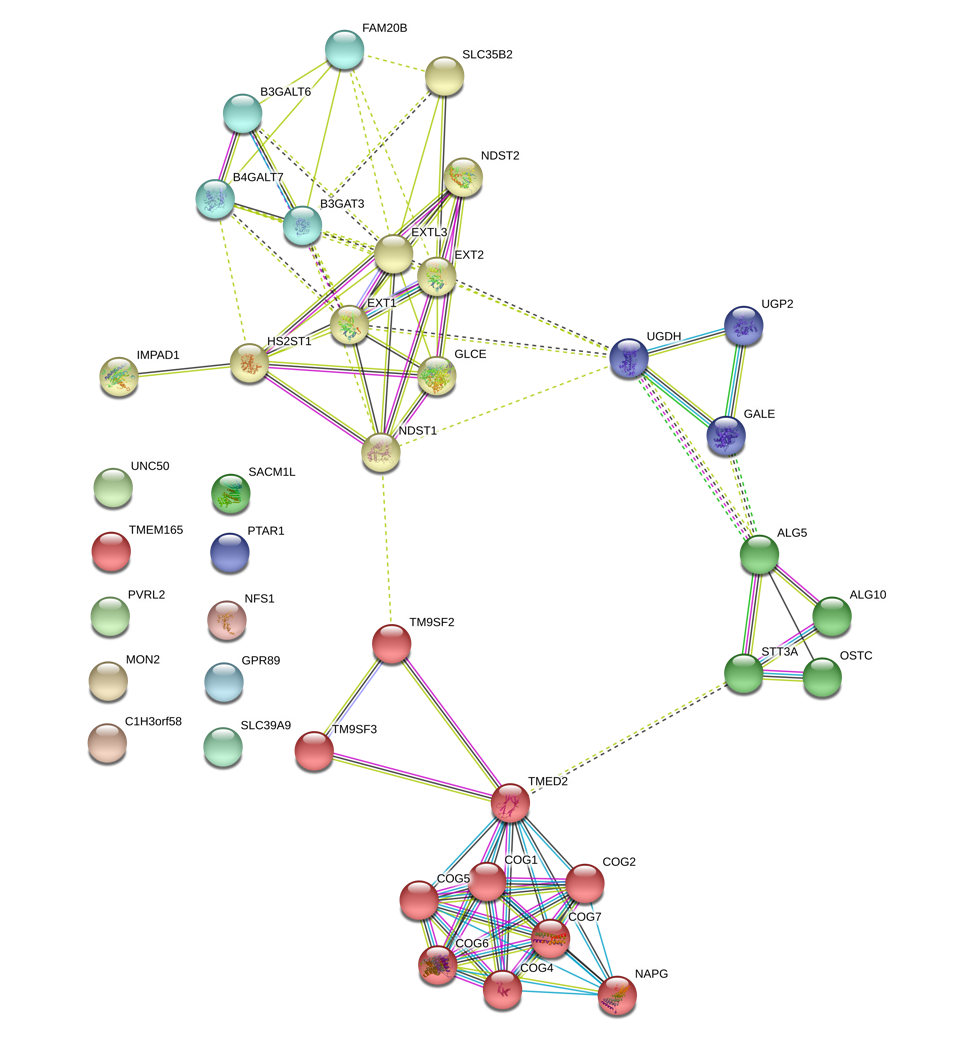

**Figure S1. STRING analysis of all candidates with suspected roles in cell entry.**

Average Plaque Number

| Concentration (microgram per ml) | Heparin | Chondroiten sulphate |
| --- | --- | --- |
| 0ug/ml | 32.4286 | 19 |
| 2.5 ug/ml | 10 | 18.3333 |
| 5ug/ml | 9 | 15.3333 |
| 10ug/ml | 4.42857 | 15.6667 |
| 100ug/ml | 1.85714 | 18.3333 |
| 200ug/ml | 3 | 18 |

**Figure S2. Virus infection blocking assays using Heparin and Chondroitin Sulfate.** Cells were treated with specified concentrations of Heparin or Chondroiten Sulfate prior to infection by BHV-1. Total viral samples were harvested and titrated on wt MDBKs cells (n=3 for HS blocking assay and n=2 for CS blocking).

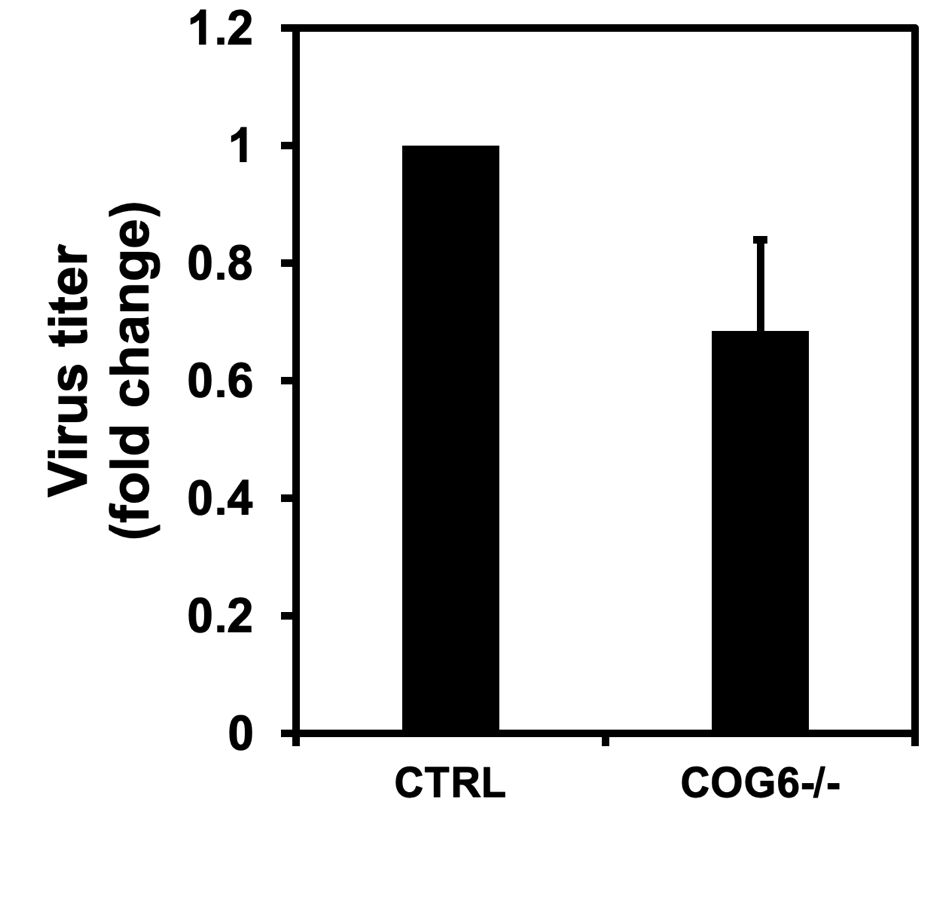

**Fig. S3. COG6KO reduced AIHV-1 replication.** Plaque assay was conducted in Cas9+/+ control cells and COG6 KO cells infected with AIHV-1. Virus titers are presented as relative numbers with that in controls cells set as 1 (n=3, p=0.016).

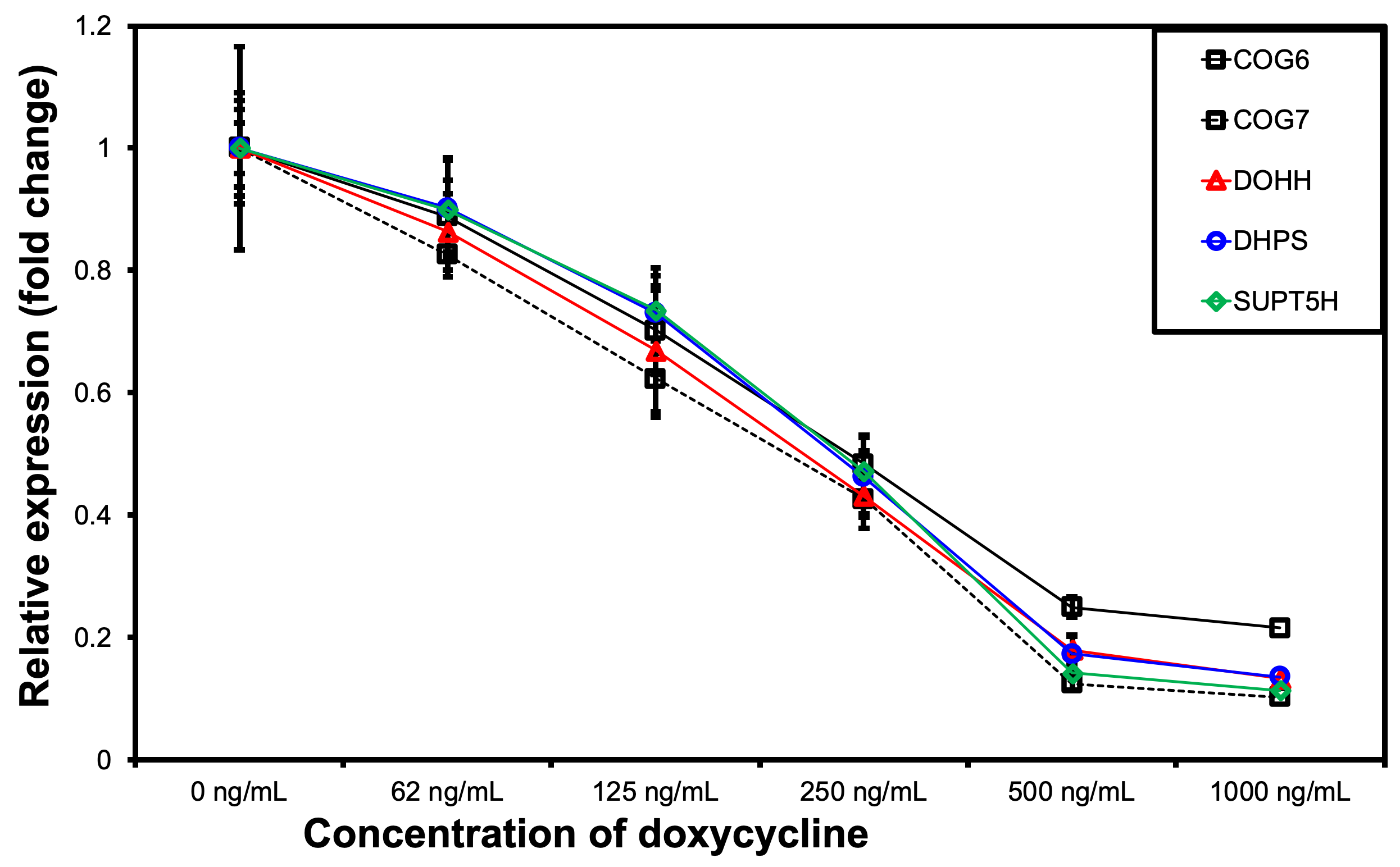

**Figure S4. Doxycycline concentration test.** dCas9 +/+ cells stably integrated with the most efficient guide from **Figure 6** for each gene were cultured in total growth media containing specified concentrations of Doxycycline 48 hours prior to mRNA sample collection and RT-qPCR (n=3 with three technical repeats for each gene).

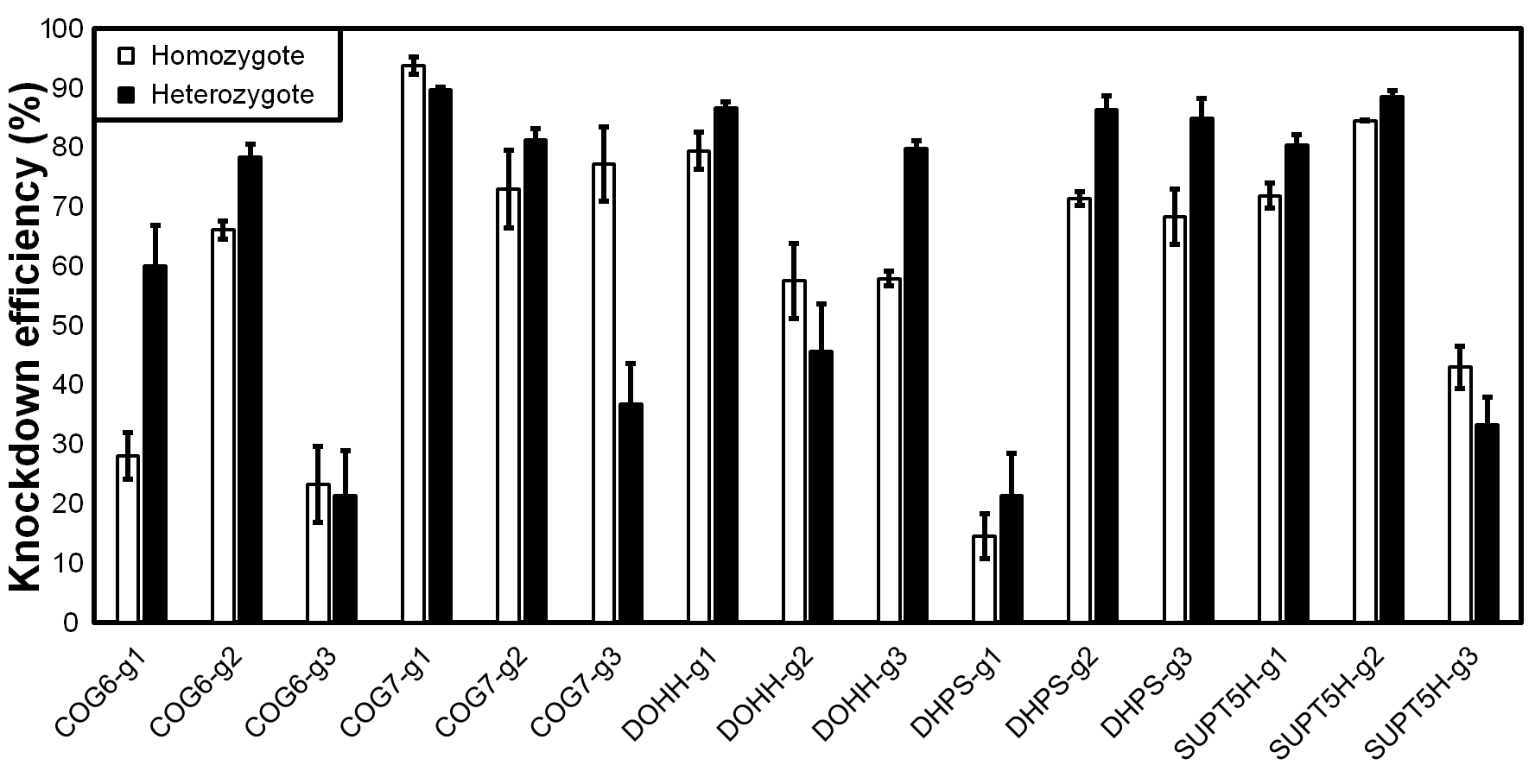

**Figure S5. Comparison of CRISPRi KD efficiency in heterozygote and homozygote dCas9 cell lines.** KD efficiency was assayed using RT-qPCR using mRNA samples collected from dCas9+/+ or dCas9/wt MDBKs transfected with PiggyBac vector expressing one of the three guides (g1,g2,g3) targeting one of the five genes tested. KD efficiency was calculated by 2^-ΔΔCT^ using GAPDH as an internal control.

**Table S1. Guide RNA sequences for CRISPRko and CRISPRi**

| **Gene** | **I.D.** | **Guide sequence** | **purpose** |
| --- | --- | --- | --- |
| **PVR** | NA | GCTCTTGTCACCATTAGGGA | KO |
| **PVRL2** | NA | GTCGGCGGCGACAATCTCAA | KO |
| **GLCE** | NA | CAGAGTCAAGTGCATAAGTG | KO |
| **HST2ST1** | NA | GATGAATGTGTAGCCGAGGG | KO |
| **COG6** | NA | TTACGAAAGACTTTACCGGT | KO |
| **COG7** | NA | GTGTTGGTAGAAATCGACC | KO |
| **COG6** | g1 | CGGGTTGCTGGTCTGCGCCG | CRISPRi |
| **COG6** | g2 | GGACAACGACAAGGTGAGCG | CRISPRi |
| **COG6** | g3 | AGTCGTAGTGCCCGCGACTG | CRISPRi |
| **COG7** | g1 | CGTGAAGGAGTGGATCAACG | CRISPRi |
| **COG7** | g2 | CCAAGAGGTGAACCACGCCG | CRISPRi |
| **COG7** | g3 | ACATCAGCGCGCTGATCCAG | CRISPRi |
| **DOHH** | g1 | AGTGAAGAATCAGCGCGCGG | CRISPRi |
| **DOHH** | g2 | TCCGATCTGGTAGCTCACTG | CRISPRi |
| **DOHH** | g3 | TGTGAAGGATCAGGTACTGG | CRISPRi |
| **DHPS** | g1 | CGAACTGTGCTTCAGCACGG | CRISPRi |
| **DHPS** | g2 | GTGCGCGATAGTCCACGCCA | CRISPRi |
| **DHPS** | g3 | GGTACAGCAAGTCAACGCCA | CRISPRi |
| **SUPT5H** | g1 | CGAGGACAGCAACTTCTCCG | CRISPRi |
| **SUPT5H** | g2 | AGGCTAAAGAGTCGCAAGAG | CRISPRi |
| **SUPT5H** | g3 | CCGAACTGGGAGTTGCGCTG | CRISPRi |

**Table S2. qPCR primers for viral genes and for CRISPRi validation**

| **Gene** | **Forward** | **Reverse** |
| --- | --- | --- |
| **ICP4** | CGGAGAGCAGCGAGGACGACGG | GCTTCGATGGCGGCGGCTATGA |
| **UL23** | CTCTGCTACCCCTTCGCCCGCTACT | AGGGTGCACACGACGAGGTTGGC |
| **VP26** | GCAGATTTTGCATGTGCTAAACGCC | CGTGGTCGTAGGTCGCAAACAT |
| **e18S** | TGTGATGCCCTTAGATGTCC | TTATGACCCGCACTTACTGG |
| **GAPDH** | CTGACCTGCCGCCTGGAGAAA | GTAGAAGAGTGAGTGTCGCTGTT |
| **COG6** | GAGTTTGGAACAGCCAGAAGAAGTA | CGAGGTGTTCTTTCTCAGAAGCAGT |
| **COG7** | TGCTCCTCATTCCCAAGATGGATAG | TTCAGGGGGAGGGACATGATATACT |
| **DOHH** | TGCTCAAGCATGAGCTGGCCTACT | CCACCTCAACAACAGGGTCAGTAGA |
| **DHPS** | GTGGAGGAGGATTTCATCAAGTGCT | CACTTCACACCCTCTGTGTTCTGCT |
| **SUPT5H** | ATTGCCTACCAGTTCACAGACACGC | ATCCGTCATCTCCTTGATAGGCACC |

**Table S3. Genotypes of CRISPR KO clones**

| **Gene** | **Clone I.D.** | **Genotype** |
| --- | --- | --- |
| PVR | 23 | -50/-50 |
| PVR | 25 | -2/-11 |
| PVR | 29 | -8/-8 |
| PVR | 32 | -5/-33 |
| PVRL2 | A4 | -1/+1 |
| PVRL2 | A11 | -4/+1 |
| PVRL2 | C11 | -4/+1 |
| PVRL2 | D5 | -19/-19 |
| GLCE | A1 | -13/+1 |
| GLCE | A7 | +1/+1 |
| GLCE | B2 | -8/-8 |
| GLCE | B8 | +1/+1 |
| HST2ST1 | A5 | +1/+1 |
| HST2ST1 | C1 | -10/-11 |
| HST2ST1 | C2 | -1/+1 |
| HST2ST1 | C4 | +1/+1 |
| COG6 | A4 | -13/-13 |
| COG6 | A6 | -1/+1 |
| COG6 | A9 | -20/-20 |
| COG6 | C3 | -152/-152 |
| COG6 | C6 | +1/+1 |
| COG7 | A3 | -32/-32 |
| COG7 | C5 | -37/-37 |
| COG7 | F3 | -13/-13 |
| COG7 | F7 | -5/-5 |
